# Growth variation of an ambrosia fungus on different tree species indicates host specialization

**DOI:** 10.1101/2025.09.08.674918

**Authors:** Marcel Hugo Decker, Peter H. W. Biedermann, Lennart J.J. van de Peppel, Jon Andreja Nuotclà

## Abstract

Ambrosia beetles rely on mutualistic fungi as a food source for themselves and especially for their offspring, yet the influence of host tree species on fungal growth and specialization is not well understood. In this study, we investigated the growth performance of the ambrosia fungus *Dryadomyces montetyi*, the primary symbiont of the oak pinhole borer *Platypus cylindrus*, on semi-artificial media infused with extracts of four tree species: *Quercus robur, Fagus sylvatica, Abies alba*, and *Pseudotsuga menziesii*. Fungal growth was quantified over time using logistic growth models, and final biomass was assessed through dry weight measurements. The growth of *D. montetyi* differed significantly among the different host tree substrates. Growth on *F. sylvatica* was comparable to that on *Q. robur*; however, both conifer-derived media (*A. alba* and *P. menziesii*) exhibited significantly reduced surface expansion and biomass accumulation. Tissue density measurements further indicated that the mycelium grown on *Q. robur* was denser than that grown on conifer media, although overall density was highest on a nutrient-rich medium without tree extract. These results demonstrate that *D. montetyi* performs best on angiosperm hosts, particularly oak, which aligns with the known host preferences of *P. cylindrus*. Our findings suggest that fungal performance is not solely determined by the typical polyphagy of ambrosia beetles but also reflects the host-related specialization of the fungal symbiont. This specialization certainly determines the host selection of *P. cylindrus* and the evolution of the tripartite interactions between this beetle, *D. montetyi*, and oak trees.

## INTRODUCTION

Ambrosia beetles are not a taxonomic classification but rather a group of beetles that independently evolved fungus-farming in the weevil subfamilies Scolytinae and Platypodinae (Farrell et al., 2001), as well as in the superfamily Lymexyloidea (Wheeler, 1986), sharing a common ecological lifestyle (Hulcr et al., 2015). They maintain an obligate mutualistic association with so-called ambrosia fungi, which spores they carry in highly specialized organs (i.e., the mycetangia (Francke-Grosmann, 1956b)) and inoculate into the wood while excavating breeding tunnels (Francke-Grosmann, 1956a; Batra, 1963, 1966). Both larvae and adults obligately rely on these fungi for nutrition (Batra, 1966). Inside their galleries, ambrosia beetles actively manage their fungal symbionts and prevent the spread of competing microorganisms (Batra, 1963; Nuotclà et al., 2019, 2021; Skelton et al., 2020; Diehl et al., 2022). Most ambrosia fungi are members of the phylum Ascomycota, particularly within the orders Ophiostomatales and Microascales and occur in genera such as *Ambrosiella, Raffaelea, Dryadomyces* and *Ophiostoma* (Massoumi et al., 2009; De Beer et al., 2022). Today, many ambrosia fungi rely entirely on beetles for survival, whereas their ancestors lived freely and spread with the help of arthropods (Mayers et al., 2015, 2015; Ranger et al., 2018).

The ambrosia beetle *Platypus cylindrus* (Fabricius, 1792; Coleoptera: Curculionidae: Platypodinae), also known as the oak pinhole borer (Baker, 1963), belongs to the Coleopteran subfamily Platypodinae (Vanderpool et al., 2018), which comprises more than 1,400 described species. The majority of these species are native to tropical regions, and with only two known exceptions, all are ambrosia beetles (Jordal, 2014). Next to *P. cylindrus*, only one other platypodine species occurs in southern Europe: *Platypus oxyurus* Dufour. It occurs in the Pyrenees, Turkey, Corsica, Calabria, and Greece and is strictly associated with silver fir (*Abies alba* Mill.) (Balachowsky, 1949). In contrast, *P. cylindrus* has been recorded on various broadleaf tree species. The principal hosts are *Quercus* spp., along with *Fagus sylvatica* L. (Strohmeyer, 1906). Additional host species include *Castanea sativa* L. (Nördlinger, 1884), *Ulmus* spp. (Strohmeyer, 1907), *Prunus avium* L. (Cassier et al., 1996), *Juglans regia* L. and *Fraxinus excelsior* L. (FNR, 2022). This diversity demonstrates the wide potential host range of *P. cylindrus* among hardwood tree species, even though oak species are preferred. This polyphagous nature of *P. cylindrus* is assumed to be closely linked to the polyphagy of its ambrosia fungi. More generally, polyphagy is an ancestral trait and possibly an evolutionary adaptation to the high floristic diversity of tropical ecosystems, where most ambrosia beetle taxa originate. In such ecosystems, narrow tree host specialization would be disadvantageous for both beetles and their fungal mutualists (Beaver, 1979, 1989).

The ecology of *P. cylindrus* has been extensively studied before the mid-20^th^ century (Knotek, 1896; Strohmeyer, 1906, 1907; Groschke, 1952; Husson, 1955; Baker, 1963; reviewed in Decker and Biedermann, in Review) while recent studies have focused on the fungi associated with *P. cylindrus* (Baker, 1963; Inácio et al., 2008, 2011; Inacio et al., 2012; Amoura et al., 2021). However, platypodine ambrosia beetles are typically associated with a richer fungus community than other ambrosia beetle groups (e.g. compared to *Xylosandrus* ambrosia beetles; Mayers et al., 2015), so the role and importance of individual microbial species for the beetles is not fully clear. The currently increasing economic relevance of *P. cylindrus* calls for a good understanding of the associated microbial players (Hulcr and Dunn, 2011; Decker and Biedermann, in Review).

An early study reported *Raffaelea ambrosiae* Arx & Hennebert as the nutritional symbiont of

*P. cylindrus* from Great Britain (Baker, 1963; Arx and Hennebert, 1965). More recent studies from Portugal and Algeria also report *Dryadomyces montetyi* (M. Morelet) M. Procter & Z.W. de Beer (synonym *Raffaelea montetyi* M. Morelet) and *Raffaelea canadensis* L. R. Batra as primary symbionts (Inácio et al., 2008, 2011; Amoura et al., 2021). However, while *P. oxyurus* shows a strict host preference for silver fir, *P. cylindrus* has been observed on several broadleaved tree species, indicating a specialization on broadleaves rather than on a single host. This strong host preference may be attributed to differences in the establishment success of their fungal symbionts. For successful colonization, nutritional fungi must quickly establish within freshly excavated tunnels and maintain a competitive advantage over other microbial taxa.

Experimental studies have shown that ethanol can act as a selective factor, enhancing the initial growth and competitive success of ambrosia fungi after breeding tunnels are inoculated with their spores (Ranger et al., 2018; Lehenberger et al., 2021). Furthermore, optimal temperature conditions (Nakashima et al., 1987) and the concentration of key nutrients such as nitrogen (Roeper and French, 1981) significantly affect fungal growth. While these factors have been extensively investigated, the role of the host tree species in shaping the growth performance of ambrosia fungi remains poorly understood. We hypothesize that the host tree species significantly affects the growth of *D. montetyi*, because of the tree-host preference of its beetle associate *P. cylindrus*, which may reflect patterns of host-tree specialization by the fungus. This would imply that co-evolution may have occurred not only between this beetle and its main fungal symbiont, but also between *D. montetyi* and oak trees. To test this hypothesis, we measured the performance of *D. montetyi* on semi-artificial growth medium infused with wood extracts from four different tree species.

## MATERIALS AND METHODS

### Fungus strain

For all our experiments we used a single strain of *D. montetyi* (F80052) from our inhouse culture collection at the chair of Forest Entomology and Protection in Stegen, Germany. This strain was isolated from *P. cylindrus* infesting an oak stump at a forest edge east of Freiburg, Germany (360 m a.s.l.) in June 2023 (see Appendix for detailed methods). A partial sequence of the ribosomal 28S large subunit (LSU) sequence of this strain is available on NCBI GenBank under the accession number PX240738.

### Fungal growth on different modified wood substrates

Wood discs of *Abies alba* Mill., *Pseudotsuga menziesii* (Mirbel) Franco, *Fagus sylvatica* L., and *Quercus robur* L. (48–56 cm diameter, ∼10 cm thickness) were obtained from local sawmills. After five days of storage at room temperature (25 °C), the discs were planed and the resulting wood chips subsequently ground with a rotor mill (18,000 rpm, 2 mm sieve). From 200 g sawdust per tree species, a 20 % stock solution was prepared by boiling it in 1 l water for 20 min at 100 °C. It was then filtered and further processed into a standardized Master-Mix medium (see Figure 1). After autoclaving, the medium was poured into Petri dishes (90 mm × 15 mm; 20 ml per dish).

**Figure 1.**
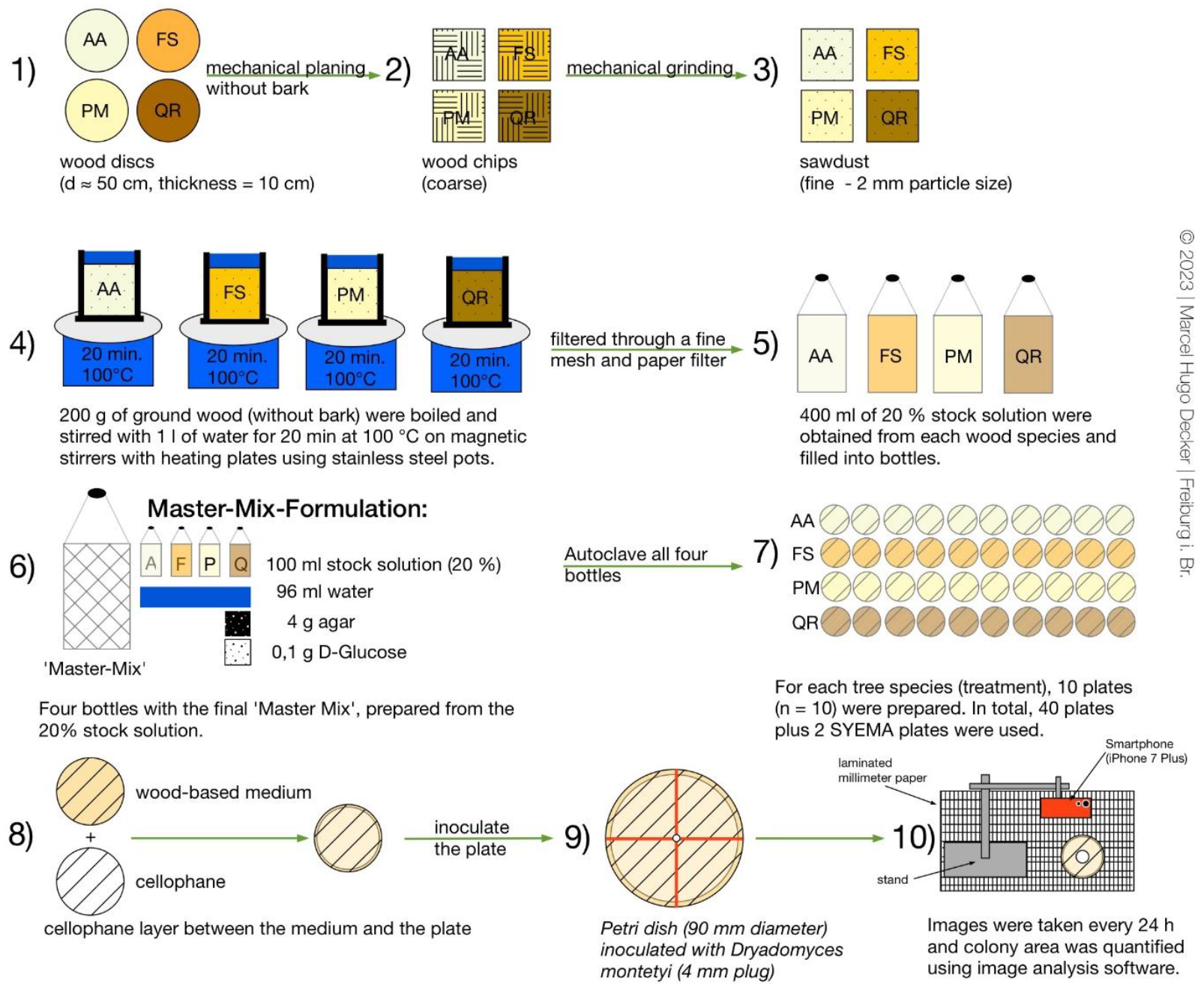
Preparation of wood-extract “Master-Mix” medium and plate assay. Wood from four host trees was processed to sawdust, extracted to 20% stocks, combined into a standardized agar medium, poured, and centrally inoculated with *D. montetyi* for growth and biomass measurements. AA= *Abies alba*, FS= *Fagus sylvatica*, PM= *Pseudotsuga menziesii*, QR= *Quercus robur*

Each plate was lined with a cellophane layer and inoculated with a mycelium plug of 2mm radius taken from a 10-day-old *D. montetyi* culture grown on nutrient rich SYEMA medium (3 g yeast extract, 10 g malt extract, 15 g agar, 100mg Streptomycin, 1 l water). Note that streptomycin is principally not needed for the current experiment but was retained to follow our usual laboratory protocol where it helps reduce bacterial contamination risk. Inoculation was performed centrally using a punch tool, with millimeter paper as positional reference. In total, 40 plates were prepared for the main experiment (10 replicates per tree species) plus two additional SYEMA plates as high nutrient reference. Fungal growth was documented every 24 h under sterile conditions by orthogonal photographs and quantified with ImageJ (version 1.53k) until either the surface of the Petri dish was fully covered, or four days had elapsed. Subsequently, the mycelium was scraped off the cellophane, dried for 5 days at 50 °C, and weighed with a precision balance (Kern PNJ, accuracy 1 mg). Biomass was calculated by differential weighing of empty versus mycelium-containing micro reaction tubes.

### Statistical analysis

First, we characterized the growth of fungal propagules over time fitting a logistic growth model to the measured fungus area data: 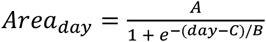

***A*** is the asymptotic maximum area, thus the theoretical maximum growth area

***B*** is the time scaling parameter, indicating growth speed

***C*** is the inflection point, thus the timepoint with fastest fungal growth

***Peak growth*** (=absolute growth at the inflexion point) has a more intuitive biological meaning than *B* and was thus calculated from the derivative 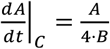.

Logistic models were fitted individually for each plate using nlsList in the statistics software R, allowing for plate-specific parameter estimates (see S-Figure 1 for individual curve fits). Standard errors (SE) were extracted to account for differences in measurement precision. We subsequently fitted weighted linear models for peak growth and the logistic growth parameters *A* and *C* to assess fungal growth across media derived from different host tree species. Each model had the form: 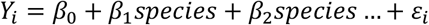,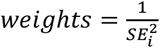 , where *Y*_i_ is the parameter of interest, β_0_ represents growth medium derived from *Q. robur* as the reference, and β_*j*_ are differences between medium obtained from *Q. robur* and each of the other three tree species as well as complete nutrient rich SYEMA medium. Post-hoc pairwise comparisons were not performed, as *Q. robur* is the main host and provides the biologically relevant baseline.

Finally, we calculated the fungal tissue density for each plate by dividing the dry weight by the total growth area on the last day of measurement. Those values where then also compared between the different growth media using linear models in the statistic software R. The *P. menziesii* treatment was excluded from this analysis because the mycelial mass was too low to be accurately measured with our method.

All analyses were conducted in R studio (Posit team, 2024) with R version 4.4.2 (R Core Team, 2024), using the nlme (Pinheiro and Bates, 2000; Pinheiro et al., 2024), dplyr (Wickham et al., 2023), ggplot2 (Wickham, 2016), writexl (Ooms, 2025) and ggbreak (Shuangbin Xu et al., 2021) packages.

## RESULTS

Fungal growth and biomass differed significantly among growth media, with *Q. robur* growth medium serving as a biologically relevant reference. The calculated maximum area (*A*) was highest on *Q. robur* (6888 mm^2^), and significantly lower on *A. alba* (by −2838 mm^2^, *p* < 0.001) and *P. menziesii* medium (by −5953 mm^2^, *p* < 0.001), whereas *F. sylvatica* (by −703 mm^2^, *p* = 0.104) and the complete *SYEMA medium* (by −411 mm^2^, *p* = 0.30) did not differ significantly from the reference (Table 1, Figure 2, see S-Table 1 for detailed model outputs).

**Table 1.** Mean growth parameters of the ambrosia fungus D. montetyi across different test media. Shown are the mean maximum colony area (A, mm^2^), the mean inflection point of the growth curve (C, days), the mean peak growth rate (A/4B, mm^2^/day) at the inflection point, and the final fungus density (µg/mm^2^).

| Grow medium | Max Area<br>(*A*, mm<sup>2</sup>) | Inflection<br>point<br>(*C*, days) | Peak growth<br>(mm<sup>2</sup>/day) | Density<br>(µg /mm<sup>2</sup>) |
| --- | --- | --- | --- | --- |
| *Quercus robur* | 6888 | 2.933 | 3641 | 0.636 |
| *Fagus sylvatica* | 6185 | 3.022 | 3526 | 0.421 |
| *Abies alba* | 4050 | 3.129 | 2297 | 0.349 |
| *Pseudotsuga menziesii* | 934 | 3.045 | 398 | - |
| SYEMA (lab substrate) | 6477 | 2.589 | 4571 | 5.737 |

**Figure 2.**
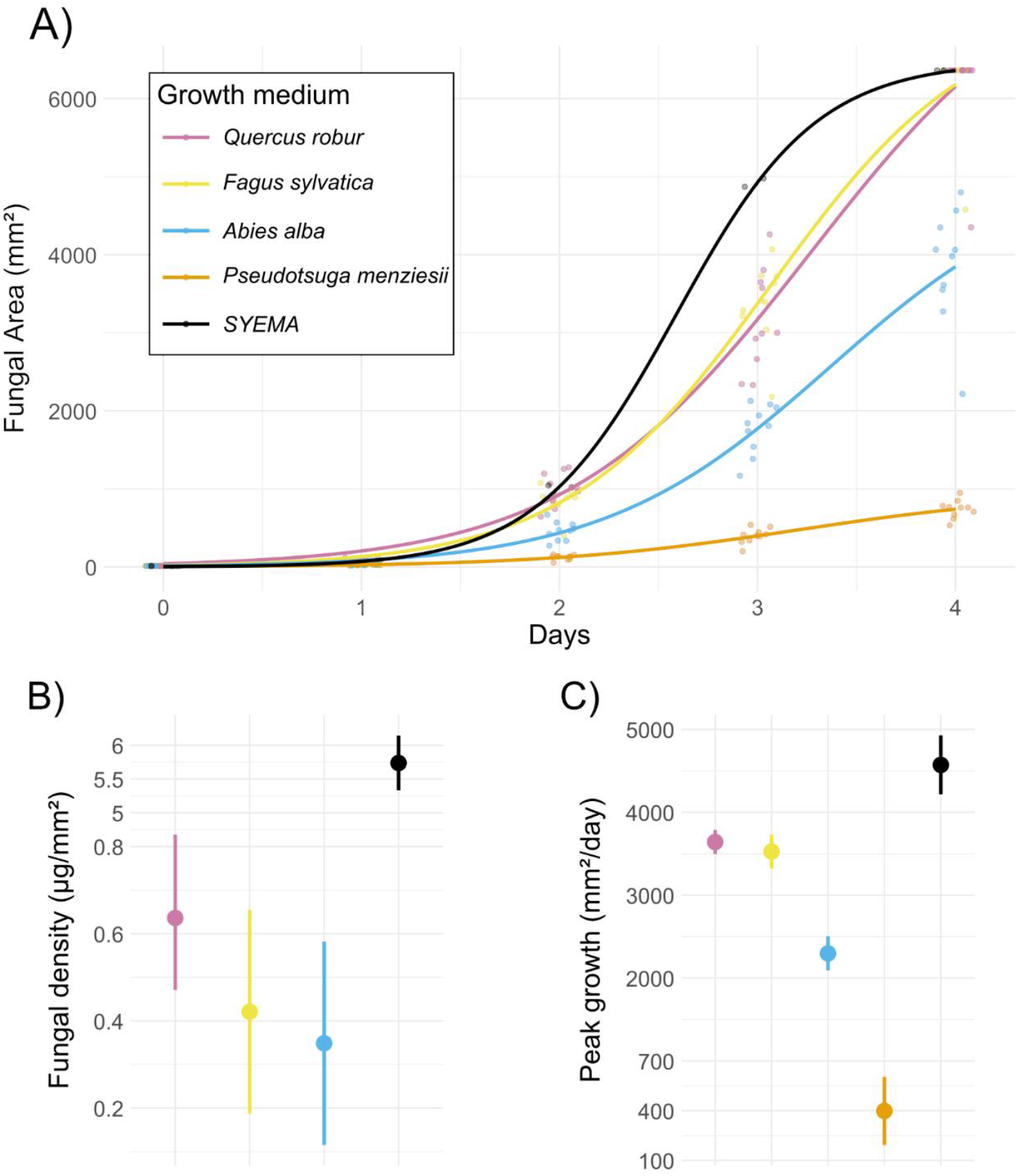
A) Logistic growth curves of fungal colony expansion on different growth media. Dots represent observed colony areas for replicate plates, with small horizontal jitter to reduce overlap, while solid lines show species-level logistic fits. B) Fungal density across media, shown as estimated means with error bars indicating standard errors. A pseudo-broken y-axis compresses the high values of SYEMA to allow clearer visualization of variation among the other media. C) Peak daily growth rates across media, shown as estimated means with standard errors. A pseudo-broken y-axis compresses the extreme values, enabling comparison of lower rates across species.

The inflection point (*C*) occurred after 2.93 days or 70 hours on *Q. robur* medium, whereas fungus on the complete *SYEMA* medium reached peak growth earlier (by −0.344 days or – 8 hours, *p* < 0.001) and *A. alba* medium slightly later (by +0.196 days or +5 hours, *p* = 0.050). C did not differ significantly between *Q. robur* and *F. sylvatica* or *P. menziesii* respectively (Table 1, Figure 2, see S-Table 1 for detailed model outputs).

Peak growth rate was 3641 mm^2^ per day in *Q. rubor* and was not significantly different in *F. sylvatica* medium (by +115 mm^2^ per day, P=0.579). Both, media sourced from *A. alba* (by - 1344 mm^2^ per day, p<0.001) and *P. menziesii* (by -3243 mm^2^ per day, p<0.001) had significantly lower peak fungal growth rates, however, on complete SYEMA medium, the peak growth rate was significantly higher than in *Q. robur* (by +931 mm^2^ per day, P=0,013). (Table 1, Figure 2, see S-Table 1 for detailed model outputs).

Tissue density was 0.636 μg /mm^2^ on *Q. robur* medium and significantly higher on complete *SYEMA* medium (by +5.101 μg /mm^2^, *p* < 0.001). We found no statistically significant fungal tissue density difference between *Q. robur* and both, *F. sylvatica* and *A. alba* respectively. (Table 1, Figure 2, see S-Table 1 for detailed model outputs).

## DISCUSSION

In summary, of all the tree infused media, the ambrosia fungus *D. montetyi* grew best on the media derived from *Q. robur* wood. However, the nutrient rich standard laboratory medium SYEMA fostered both significantly higher growth rates and much higher fungal tissue density.

### Methodology

We demonstrate that growing *D. montetyi* on a semi-artificial medium composed of wood extract and agar is an effective method to differentiate fungal performance on different host tree species, as we could clearly observe significant differences in radial growth parameters and mycelial density between treatments. We also show that radial growth or fungal surface area alone is not sufficient to measure fungal performance, as two colonies with a similar final growth area, like *Q. robur* and the nutrient rich SYEMA medium can produce mycelium that strongly differs in its density. Although we can recommend density calculated from dry mass and final growth area as a reliable metric (see e.g. Lehenberger et al., 2021), some caution is advised. The mycelial mass is generally only in the order of a few milligrams when fully covering standard sized petri dishes. Thus, very high precision measurements are needed to accurately measure slow growing fungi. In our study, we had to discard the measurements for the *P. menziesii* medium, as the fungal growth was below the detection threshold of our one-milligram precision scale. The amount of biomass produced on the other types of media was also low, but measurable, as the fungus covered almost the entire plate after three to four days. To obtain sufficient fungal biomass for treatments with slow growing fungi, we recommend future studies to extend the duration of the experiment and use larger petri dishes or race tubes (e.g. Ryan et al., 1943). This allows prolonged fungal growth and a better comparison of growth characteristics among the treatments with faster growing fungi.

We show that *D. montetyi* performs significantly better on the host tree species that are preferred by the host beetle *P. cylindrus*. Not only does it grow faster on *Q. robur* and *F. sylvatica* medium in comparison to the medium containing extracts of the two gymnosperm species, but it covers significantly larger areas before the growth speed declines. Additionally, it did produce denser mycelium on *Q. robur* relative to the others, although these differences were not statistically significant because of relatively large standard errors (see Figure 2).

Our experiment is limited to showing differences in fungal growth but cannot explain the cause of these differences. The reduced fungal growth on the *P. menziesii* and *A. alba* medium could be because of inhibition caused by defensive chemicals produced by the host tree, differences in sugar concentration or the lack of certain essential micronutrients. To find an explanation for the observed differences in fungal growth between the different treatments, future studies need to analyze the wood extracts and determine the concentration of macro and micronutrients and tree defensive chemicals before and after the fungus has grown on the plates.

### Tree host specialization of *D. montetyi*

Interestingly, *D. montetyi* is not strictly associated with *P. cylindrus*, but has also been found in association with *Xyleborus dryographus* and *Xyleborus monographus* that also inhabit oak trunks (Gebhardt et al., 2004). The *D. montetyi* strains associated with these three different beetle species do not show significant genetic variation, indicating recent and potentially repeated horizontal transfer. Horizontal transfer between these beetle species has been attributed to their shared use of oak trunks and to the fungus’s ability to invade neighboring galleries through woody tissue, a process which should be facilitated by its faster growth rate relative to other ambrosia fungi (Gebhardt et al., 2004). An alternative explanation could be joint usage of tunnels as has been recently reported to be quite common in some ambrosia beetle species (Scabbio et al., in review). Occasional horizontal symbiont transfer between *P. cylindrus* and other unrelated ambrosia beetle species may have prevented the evolution of a strong host-symbiont fidelity of *D. montetyi*.

As most ambrosia beetles have a broad range of host trees, we did not expect that their fungi are more selective (Beaver, 1979, 1989). Our findings do not support a broad potential host range as we observe that *D. montetyi* grows significantly faster on angiosperm compared to gymnosperm tree-hosts, indicating that there is at least some degree of specialization by the fungus. Unfortunately, there are very few experimental studies that further tested this hypothesis. Castrillo et al. (2012) found significant differences in radial growth of different strains of the fungal symbiont of *Xylosandrus germanus* (Blandford, 1894) on medium infused with sawdust of different host tree species. However, they only measured radial growth once after two days and did not measure fungal biomass or density. Therefore, it remains unclear whether the case of *D. montetyi* is an exception or whether host tree specialization occurs more frequently.

It is not known if host tree specialization of *D. montetyi* also affects host beetle fitness or behavior and thus may select for host tree preference by the beetle. Alternatively, in evolutionary history, specialization by beetle hosts may have resulted in subsequent specialization by the fungus. Some studies have experimentally measured ambrosia beetle fitness on semi-artificial substrates using sawdust of different host tree species and found conflicting results. A study on *Xyleborus glabratus* (Eichhoff, 1877) found lower success rates and decreased productivity on non-preferred host trees (Maner et al., 2013) and a study on *X. germanus* found significant differences in progeny produced among four different host tree species even though this beetle is regarded to be a host generalist (Castrillo et al., 2012). Contrary to this, a study on the host-tree generalist, *Xyleborinus saxesenii* (Ratzeburg, 1837) found no effect of host-tree species on beetle fitness (Melet and Biedermann, 2024).

Our findings may indicate why studies come up with opposing results, as we report significant improvements of fungal growth speed and density for the very nutrient rich SYEMA vs *Q. robur* medium. This highlights the importance of considering the nutrient content of artificial media that have been used in all the studies above, as it is almost impossible to measure fungal biomass directly on the wood due to the potential ability of the fungus to penetrate the wood (Mathiesen-Käärik, 1953; Francke-Grosmann, 1956a). For future studies it would be interesting to test if the differences in fungal growth rate that we observe also align with difference in fitness of *P. cylindrus*, however, attempts to rear *P. cylindrus* on artificial medium have yet not been successful (Decker, 2023).

To understand the complex interactions that lead to host-symbiont fidelity it is crucial to study all partners involved. Except for a few studies mentioned above, most studies have focused on the host tree preference of the beetle (Castrillo et al., 2012; Maner et al., 2013; Melet and Biedermann, 2024) but have not considered possible host tree specialization by their symbiotic fungi. Testing both the performance of ambrosia beetles and their fungal symbionts on different tree hosts will help in identifying and understanding the drivers that shape host-fidelity and the evolution of obligate fungus-farming.

## AUTHOR CONTRIBUTIONS

**MHD:** Conceptualization, Data curation, Investigation, Visualization, Writing - original draft, Writing - review & editing

**PHWB:** Conceptualization, Resources, Supervision, Writing - review & editing

**LJJvdP:** Conceptualization, Supervision, Writing - original draft, Writing - review & editing

**JAN:** Supervision, Formal analysis, Writing - original draft, Writing - review & editing

## FUNDING

LJJvdP was supported by the Netherlands Organization for Scientific Research (NWO-Rubicon grant no. 019.211EN.031) and an Eva Mayr-Stihl Fellowship. MHD is supported by a scholarship from the German Academic Scholarship Foundation (Studienstiftung des deutschen Volkes). We acknowledge support from the Open Access Publication Fund of the University of Freiburg.

## ACKNOWLEDGMENTS

We thank Katrin Hübscher, Andrea Andris, and Hannes Freitag for their assistance in the laboratory, and Angela Haury for support in the office.

## CONFLICT OF INTEREST

The authors declare that the research was conducted in the absence of any commercial or financial relationships that could be construed as a potential conflict of interest.

## Notes

### Competing Interest Statement

The authors have declared no competing interest.

