## Supplementary Material for "Growth variation of an ambrosia fungus on different tree species indicates host specialization"

### Fungal isolation from *P. cylindrus* via crush method

Male and female imagines, pupae, larvae, as well as material from galleries were processed with this method. One individual or substrate sample was mechanically crushed in 100 µl PBS in a sterile micro reaction tube and then suspended in an additional 900 µl PBS (stock solution, c = 1). A dilution series was prepared up to 1:1000 (c = 0.001); 100 µl of each dilution were spread on SYEMA plates. For each object, four plates were prepared and incubated and sub cultured as described in 3.2.1. From each plate, ten fungal colonies were selected and transferred to new plates, resulting in up to 40 plates per object and dilution series. Overall, ten female beetles, five male beetles, three larvae, three pupae, and one gallery sample were analyzed with this method.

### Direct inoculation from gallery substrate

Small pieces of mycelium were scraped directly from the surface of fresh beetle galleries with sterile needles and inoculated into SYEMA plates at up to three points per plate, maximally distant from each other. In total, three gallery samples were analyzed.

### DNA determination of fungi

Seven days after incubation at 25 °C in darkness, isolates were grouped according to their morphology, and two representatives per morphotype were selected and recultivated on SYEMA plates with a cellophane overlay. DNA was extracted from the mycelium either following Izumitsu et al., (2012) or using the ZymoBIOMICS DNA Miniprep Kit (with additional pretreatment by freezing in liquid nitrogen for 5 s and mechanical grinding). PCR amplification was performed with 1 µl DNA, the Q5® High-Fidelity 2X Master Mix, the forward primer LR0R (Vilgalys and Hester, 1990) and the reverse primer JH-LSU-369rc (Li et al., 2016). Thermal cycling conditions were: initial denaturation at 95 °C for 3 min; 35 cycles of (a) 30 s at 95 °C, (b) 30 s at 52 °C, (c) 30 s at 72 °C; followed by a final elongation step at 72 °C for 10 min. PCR products were visualized by 1% agarose gel electrophoresis (40 ml TAE buffer; 0.6 g agarose; GelRed as stain) using a Vilber E-Box-CX5.TS-Edge system, purified with the PCR Purification Kit (Jena Bioscience), and sequenced in both directions by Sanger sequencing (StarSEQ, Mainz). The obtained nucleotide sequences were edited and analyzed using Geneious Prime software (Dotmatics).

### Supplementary table and figure

S-Table 1 Results of weighted linear models testing differences in fungal growth parameters across growth media, using *Quercus robur* as the reference. Plate-level logistic parameters (A, B and C) were first estimated for each replicate according to $Area_{day} = \frac{A}{1 + e^{-(day-C)/B}}$ and then compared between species/media using weighted linear models with standard errors as weights. Peak growth (growth rate at the inflexion point C) was calculated using the derivative $\frac{dA}{dt}\Big|_C = \frac{A}{4 \cdot B}$. Fungus density was calculated by dividing fungal dry weight by its total growth area on the last measurement day. Shown are parameter estimates (relative to *Q. robur*), standard errors, t values, and p values. Significant differences ($p < 0.05$) indicate deviation from the reference medium.

| Parameter | Growth medium | Estimate | Std. Error | t value | p value |
|---|---|---|---|---|---|
| *A* max area | *Q. robur* (reference) | 6888 | 374 | 18.397 | |
| [mm$^2$] | *F. sylvatica* | -702 | 422 | -1.664 | 0.104 |
| | *A. alba* | -2838 | 436 | -6.508 | <0.001 |
| | *P. menziesii* | -5953 | 377 | -15.775 | <0.001 |
| | *SYEMA* | -411 | 389 | -1.056 | 0.298 |
| *C* inflexion point | *Q. robur* (reference) | 2.933 | 0.071 | 41.567 | |
| [days] | *F. sylvatica* | 0.089 | 0.080 | 1.119 | 0.270 |
| | *A. alba* | 0.196 | 0.097 | 2.028 | 0.050 |
| | *P. menziesii* | 0.112 | 0.099 | 1.135 | 0.264 |
| | *SYEMA* | -0.344 | 0.074 | -4.635 | <0.001 |
| Peak growth | *Q. robur* (reference) | 3641 | 145 | 25.058 | <0.001 |
| *A*/(4**B*) | *F. sylvatica* | -115 | 205 | -0.559 | 0.579 |
| [mm$^2$/day] | *A. alba* | -1344 | 205 | -6.540 | <0.001 |
| | *P. menziesii* | -3243 | 205 | -15.781 | <0.001 |
| | *SYEMA* | 931 | 356 | 2.615 | 0.013 |
| fungus density | *Q. robur* (reference) | 0.636 | 0.165 | 3.852 | |
| [µg/mm$^2$] | *F. sylvatica* | -0.215 | 0.233 | -0.921 | 0.365 |
| | *A. alba* | -0.287 | 0.233 | -1.231 | 0.229 |
| | *SYEMA* | 5.101 | 0.404 | 12.615 | <0.001 |

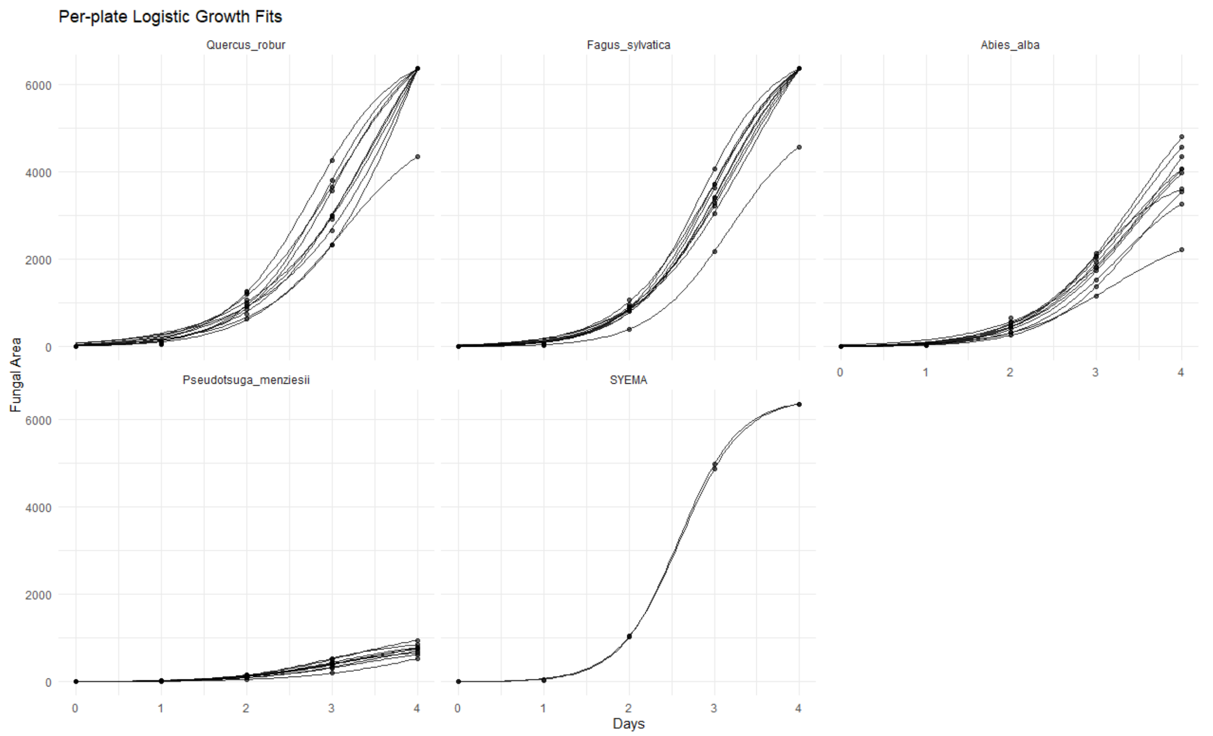


S-Figure 1 Logistic growth curves of each fungus plate observed during the experiment. Points represent observed plate-level growth measurements (Area, $mm^2$) over time (days). Lines show predicted logistic fits for each individual plate based on estimated parameters A, B and C of the curve Area= $\frac{A}{1+e^{-(day-C)/B}}$. Panels correspond to different host media.
